# Improving reproducibility in pharmacogenomic screens through cross-study benchmarking

**DOI:** 10.64898/2026.04.29.721574

**Authors:** Kristen Nader, Filipp Ianevski, Aleksandr Ianevski, Tero Aittokallio

**Affiliations:** Institute for Molecular Medicine Finland (FIMM), HiLIFE, University of Helsinki, Helsinki, Finland; iCAN Digital Precision Cancer Medicine Flagship, University of Helsinki and Helsinki University Hospital, Finland; Institute for Cancer Research, Department of Cancer Genetics, Oslo University Hospital, Oslo, Norway; Oslo Centre for Biostatistics and Epidemiology (OCBE), Faculty of Medicine, University of Oslo, Oslo, Norway

**Keywords:** Pharmacogenomics, Reproducibility, Drug-response profiling, Cancer cell models, Large-scale screening, Benchmarking

## Abstract

Drug-response measurements across pre-clinical pharmacogenomic studies remain poorly correlated, which limits biomarker discovery, precision oncology, and predictive modelling. The drivers of this inconsistency have been debated but not yet resolved. By integrating 15 pharmacogenomic studies encompassing 760 small-molecule compounds, 1,111 cell models, and 9.8 million dose-response measurements, we demonstrate that dose-response metric is the strongest driver of inconsistency, followed by experimental factors, such as treatment duration, plate format, and viability readout; in contrast, cell line molecular features contribute only minimally to reproducibility. Among drug classes, hormone therapies and PARP inhibitors show the highest concordance, whereas antimetabolites, topoisomerase inhibitors, and mitotic inhibitors exhibit substantial response variability across studies. To improve consistency, we developed a Drug Response Score (DRS), a proximity-weighted measure that emphasize pharmacologically informative concentrations near IC_50_, and we demonstrate in systematic benchmarking how DRS markedly improved cross-dataset concordance. Applications to patient-derived neuroblastoma organoids and leukemia patients’ primary cells demonstrate that DRS improves replicate-level consistency in patients’ drug-response profiles. To improve reproducible pharmacogenomic studies, we make openly available an integrated Drug Response Resource (iDRR, https://aittokallio.group/iDRR/), a standardized 15-dataset portal that supports robust biomarker discovery and cross-study benchmarking.

## Introduction

The promise of precision oncology is based on the ability to predict therapeutic responses from genomic and molecular features. Over the past decade, large-scale pharmacogenomic initiatives including the Cancer Cell Line Encyclopedia (CCLE)^1^, the Genomics of Drug Sensitivity in Cancer (GDSC)^2,3^, the Cancer Therapeutics Response Portal (CTRP)^4^, and the PRISM Repurposing portal^5^ have collectively generated millions of drug-response measurements across molecularly characterized cancer cell lines (*in vitro* models) and patient-derived living biopsies (*ex vivo* patient cells and organoids). These resources, supported by substantial public and private investment, have become central to biomarker discovery, drug repurposing, and mechanistic interrogation of cancer vulnerabilities^6,7^.

However, systematic inconsistencies in drug response measurements have raised fundamental concerns about the translational utility of *in vitro* pharmacogenomic datasets^8–14^. A landmark comparison of CCLE and GDSC datasets revealed that drug sensitivity profiles of the same cancer cell lines correlated poorly, despite well-aligned genomic profiles between the two study sites^15^. This discrepancy, widely described as a reproducibility crisis in pharmacogenomics^9,10,15,16^, undermines confidence in translational use of pre-clinical findings. The scale of investment in assay development, compound libraries, and cell line profiling is reflected in a global high-throughput screening market valued at USD 21 billion in 2023 and projected to reach USD 42 billion by 2030^17^. Combined with extensive public-private support for pharmacogenomic infrastructures^2^, the inability to generate consistent measurements represents, therefore, not only a scientific barrier but also a major source of wasted resources.

Beyond resource inefficiency, such discordance has significant scientific consequences as well. Machine learning (ML) models trained on inconsistent datasets lead to spurious biomarkers and drug-gene associations, yielding predictions that fail in prospective or clinical testing. This distorts drug prioritization, falsely promoting agents with apparent activity in single study, while overlooking more reproducible candidates^16^. Because *in vitro* and *ex vivo* pharmacogenomic studies often guide drug repurposing and prospective validation, such inconsistency propagates errors downstream, inflating costs and weakening the translational bridge between initial pharmacogenomic screening and translational applications. The challenges in AI-based discovery are further compounded by the reliance of most ML approaches on single training datasets^18,19^, rendering them especially vulnerable to dataset-specific artifacts, instead of using robust and reproducible biological signals.

The established drivers of drug response variability are diverse, including differences in culture conditions, assay platforms, curve-fitting methods, and the use of divergent summary metrics, such as half-maximal inhibitory concentration (IC_50_), area under the curve (AUC), or drug sensitivity score (DSS)^20–22^. Previous harmonization efforts have provided valuable insights by comparing two or three datasets and proposing adjusted metrics for specific analytical differences, such as variable concentration ranges or assay-specific effects^15,16,23–26^. However, these studies addressed only individual sources of variability, without systematically disentangling their relative contributions or providing generalizable solutions applicable across diverse pharmacogenomic resources. Furthermore, even if open-access pharmacogenomic databases exists^27–30^, their use in larger-scale, multi-study reproducibility analyses and benchmarking of response metrics is limited.

Here, we report the most systematic investigation of pharmacogenomic reproducibility to date, integrating 15 large-scale datasets spanning 760 compounds, 1,111 cancer models, and more than 9.8 million dose-response measurements. Using this resource, we benchmarked cross-dataset reproducibility, identified experimental, molecular and analytical factors underlying inconsistency, and developed improved methodological solutions, including consensus curve modeling and the Drug Response Score (DRS), to enhance concordance. This work presents a quantitative framework for assessing and improving reproducibility in pharmacogenomic studies, identifies the origins of cross-dataset variability, introduces approaches that improve assay reliability, as well as provides the integrated Drug Response Resource (iDRR) and evidence-based guidelines to strengthen preclinical drug response studies for biomarker discovery, drug development, and precision oncology.

## Results

### Comprehensive data harmonization across pharmacogenomic studies

We systematically curated drug-response assay annotations and data from 15 large-scale pharmacogenomic studies spanning more than a decade of cancer drug screening (**Fig. 1a**). The integrated dataset comprises 10 monotherapy dose-response studies^1–5,24,29–36^ and 5 combinatorial dose-response matrix studies^37–41^, collectively representing diverse experimental platforms, culture conditions, and cell viability readouts (**Table 1**; **Suppl. Table 1**). From the drug combination studies, single-agent dose-response measurements were extracted by analyzing drug response profiles at zero concentration of the partner drug, enabling direct comparison with dedicated single-agent assays. To ensure robustness, we retained only the drug-cell line pairs that were independently measured in at least two studies, yielding 195,317 unique pairs spanning 760 compounds and 1,111 molecularly characterized cell lines (9,846,840 individual dose-response measurements). This harmonized collection constitutes the integrated Drug Response Resource (iDRR), which is used here for all the subsequent analyses and is made publicly available to the community (https://aittokallio.group/iDRR).

**Figure 1.**
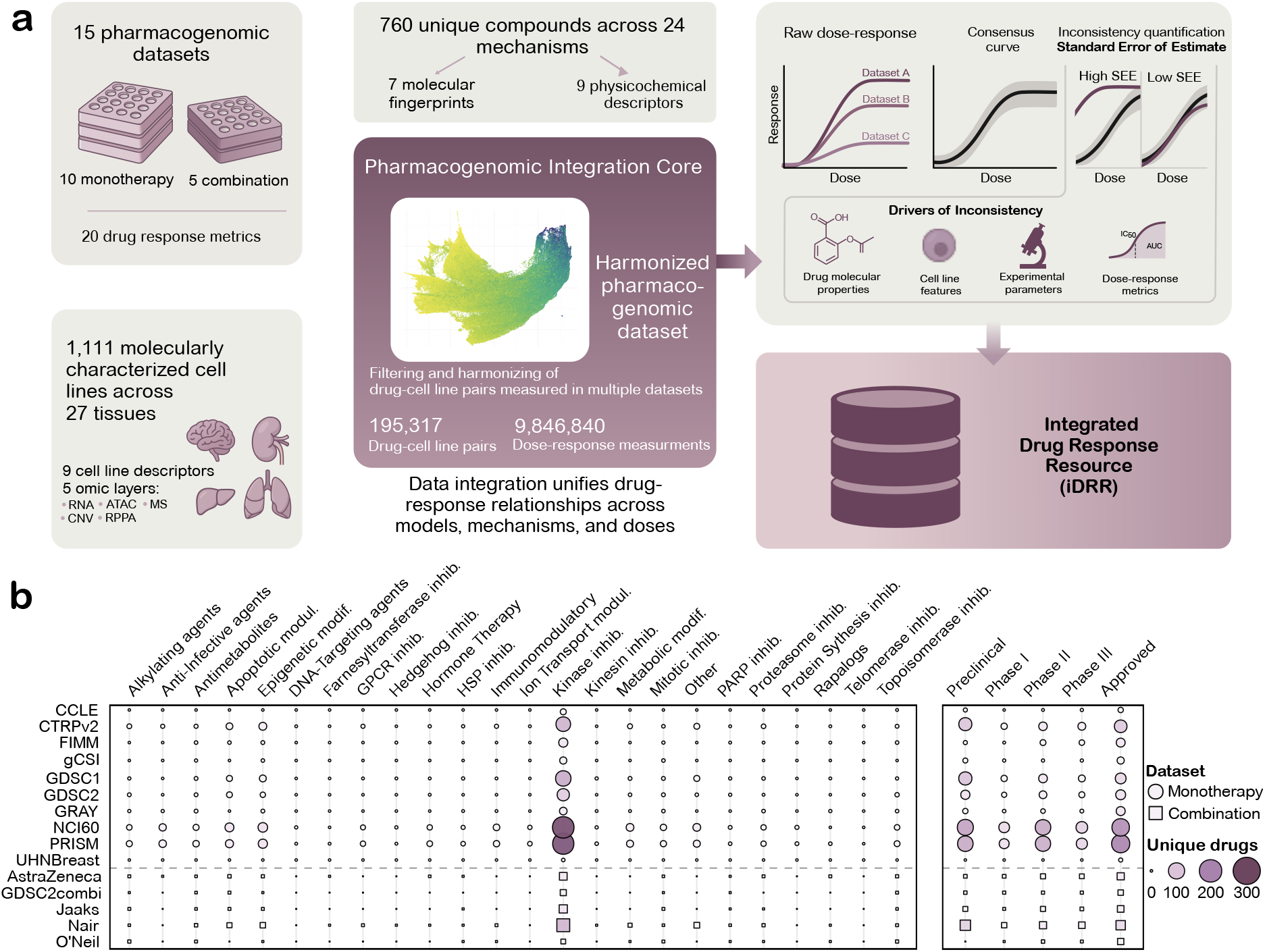
Construction and harmonization of an integrated pharmacogenomic resource. (**a**) Schematic of the pharmacogenomic integration pipeline. Drug response profiles from 15 large-scale pharmacogenomic datasets spanning 1,111 molecularly characterized cell lines and 760 compounds were harmonized using PubChem compound identifiers (CID) and Cellosaurus accession identifiers. After filtering to retain drug-cell line pairs measured in at least two datasets, all dose-response measurements for drug-cell line pairs were jointly modeled to generate consensus dose-response curves. A standard error of estimate (SEE) derived from the consensus curves quantifies cross-dataset inconsistency in the integrated Drug Response Resource (iDRR). (**b**) Bubble plot summarizes drug coverage across datasets by mechanism of action and clinical development stage. Rows represent individual pharmacogenomic studies, and columns indicate 24 therapeutic classes and clinical phases. Bubble size corresponds to the number of unique drugs within each dataset-class combination, with the inset highlighting the distribution of compounds across development stages.

**Table 1.**
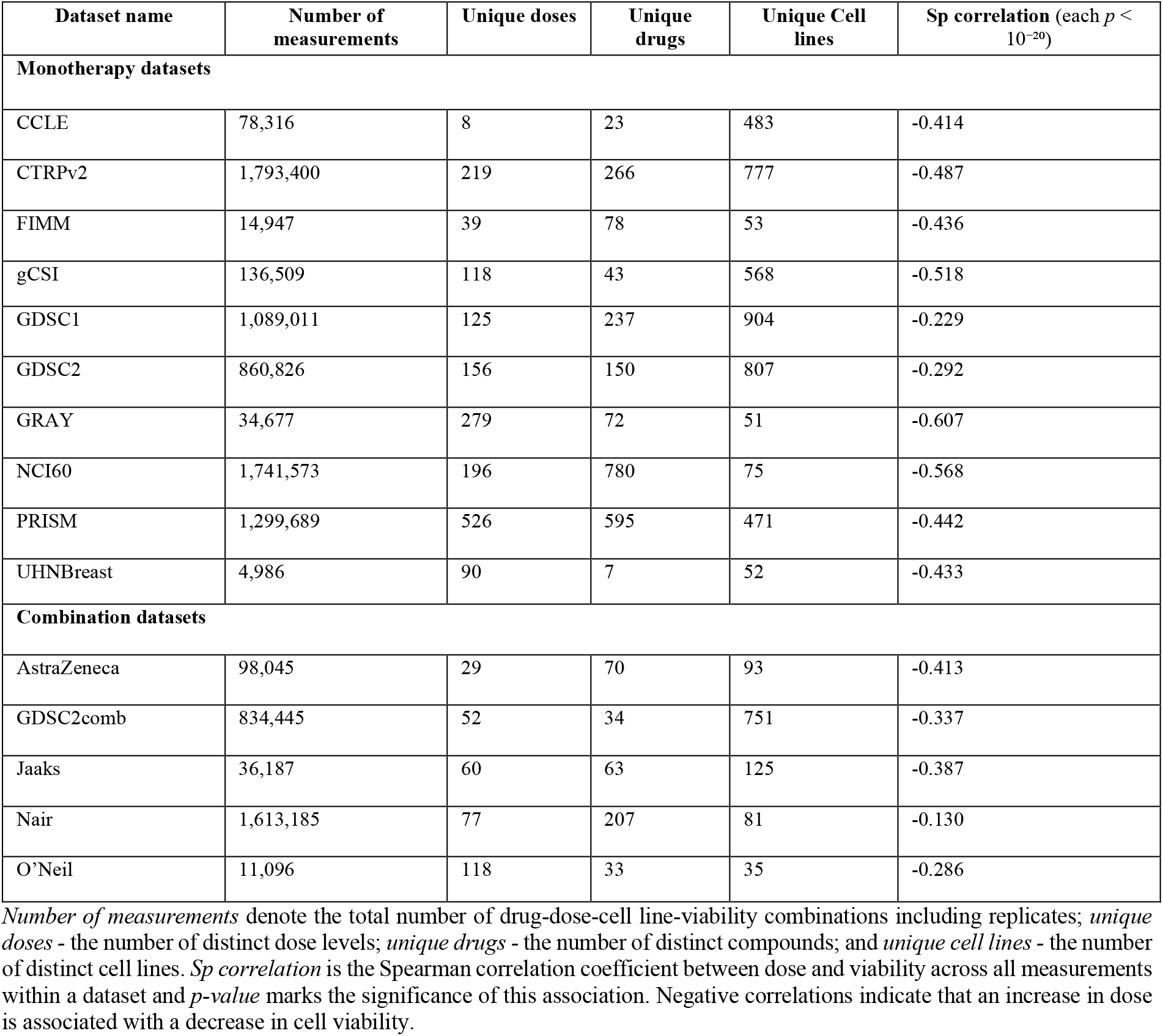
The pharmacogenomic dose-response datasets included in this study.

The size of these datasets varied by 2.5 orders of magnitude, from focused screens of a few thousand measurements (4,986 in UHNBreast) to comprehensive campaigns exceeding 1.7 million measurements (CTRPv2), with corresponding differences in the breadth of doses, compounds, and cell lines tested (**Table 1**). Notably, the strength of the global dose-viability relationship varied considerably, with Spearman correlations ranging from -0.61 (GRAY) to -0.13 (Nair). This range reflects fundamental differences in screening strategy, where datasets with stronger negative correlations typically comprise compounds pre-selected for anticipated activity, whereas weaker correlations are characteristic of large-scale discovery screens that survey chemically diverse libraries, in which most compounds are expected to show limited activity against any given cell line. Consistent with this interpretation, the proportion of dose-response curves with IC_50_ values falling within the tested concentration range was strongly associated with the global dose-viability correlation across datasets (**Suppl. Fig. 1**; Spearman ρ = 0.72, *p* = 0.00334).

The iDRR resource encompasses agents across the full spectrum of clinical development, from preclinical tool compounds to approved therapies (**Fig. 1b**). As expected, kinase inhibitors form the largest mechanistic class, followed by epigenetic modifiers, apoptotic modulators, and metabolic modifiers. Mapping the mechanisms of action and clinical phases across datasets revealed that some resources, such as PRISM, GDSC and CTRPv2, provide a broad coverage of both investigational and approved compounds, whereas others contribute towards more specialized compound subsets. This comprehensive and harmonized resource, which captures the current small-molecule landscape of cancer therapies, was then used to investigate the drug properties, cell line features, experimental parameters, and response metrics that most contribute to the drug response variability across the studies.

### Consensus dose-response modelling and quantification of inconsistency

To compare dose-response measurements across diverse assays and pharmacogenomic studies, we developed a consensus dose-response curve framework that pools all available data points for each drug-cell line pair and fits a single four parameter logistic model (4PL) to the combined measurements (**Fig. 2a**). For each contributing dataset, we quantified residuals between the observed responses and the consensus fit and summarized them as a per-dataset standard error (SE). The Standard Error of Estimate (SEE) for a given drug-cell line pair was then defined as the average SE across all contributing datasets (see **Methods**), effectively capturing how tightly the individual experiments from different datasets agree with the consensus dose-response curve.

**Figure 2.**
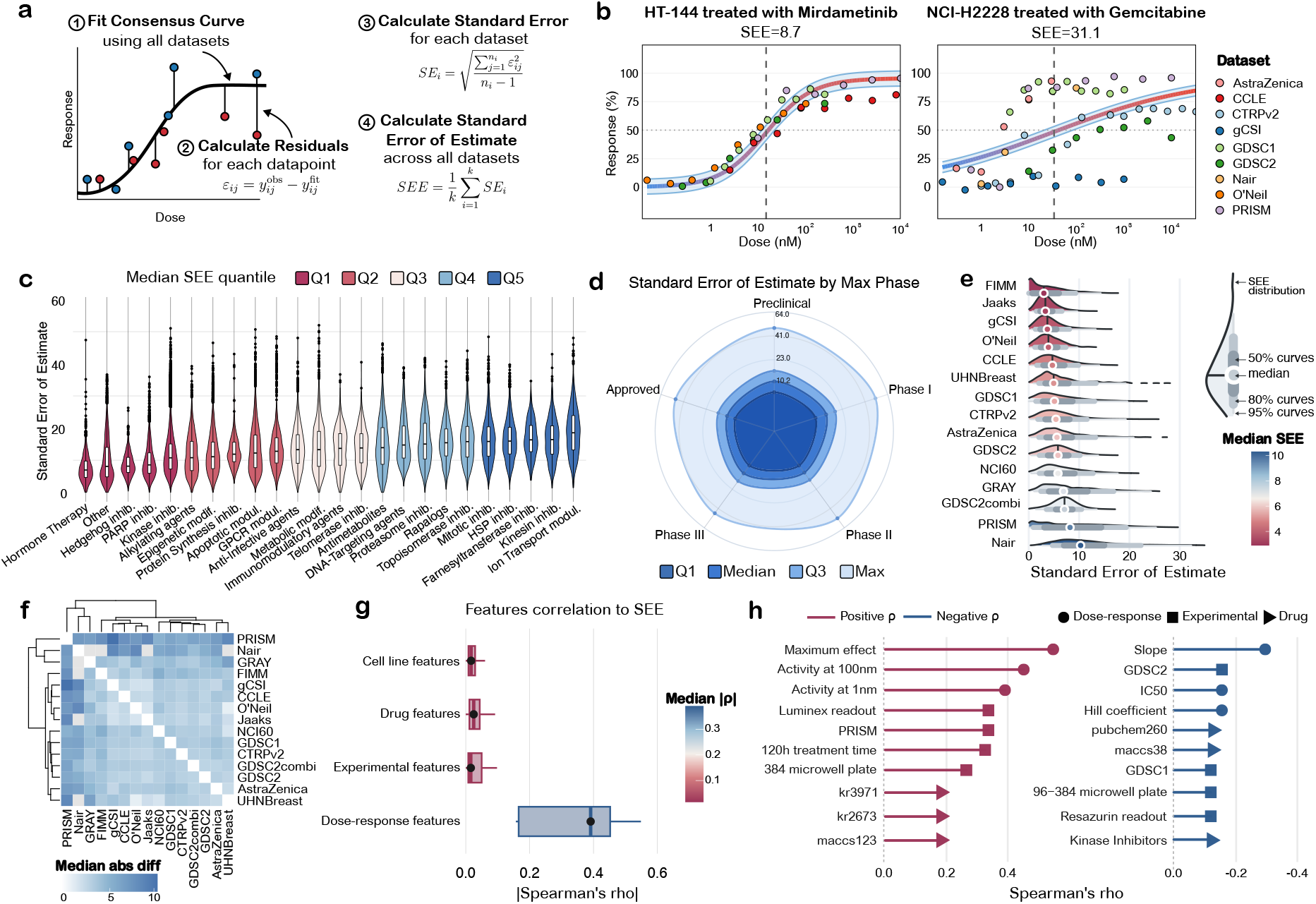
Quantifying inconsistency in drug-response measurements across datasets. (**a**) Schematic of the standard error of estimate (SEE) computation for each drug-cell line pair. A consensus dose-response curve is fitted across all datasets, residuals between observed and fitted responses are calculated, a standard error is obtained for each dataset from the squared residuals, and the SEE is defined as the average standard error across datasets. (**b**) Example drug-cell line pairs with relatively low (HT-144 treated with mirdametinib) and high (NCI-H2228 treated with gemcitabine) SEE values. Points represent measurements from individual datasets, and the blue line and band show the consensus curve and its uncertainty. (**c**) Violin plots of SEE stratified by drug mechanism of action classes, illustrating differences in cross-dataset consistency among therapeutic categories. Differences across categories were assessed using a Kruskal-Wallis test. (**d**) Radar plot summarizing SEE distributions for compounds at different stages of clinical development. (**e**) Density plots of SE for each pharmacogenomic dataset. Kruskal-Wallis test showed a significant difference across the 15 datasets (χ^2^(14) = 29,162, *n* = 477,225, *p* < 10^−16^) with a small effect size (*H* = 0.061). (**f**) Hierarchical clustering heat map of pairwise median absolute differences correlations in SEE between datasets. (**g**) Spearman correlations between SEE and four feature groups: cell-line, drug, experimental, and dose-response characteristics, showing that experimental and dose-response factors account for most of the variability in consistency. (**h**) Top individual features associated with SEE. Symbol shapes indicate feature groups (dose-response, experimental, or drug features) and line color indicates the direction of association (positive or negative Spearman correlation ρ). Although several cell-line features (clinical annotations and multi-omics measurements) were evaluated, they did not appear among the top individual correlates of inconsistency.

Example drug-cell line pairs illustrate the range of dose-response behaviours captured by the SEE metric (**Fig. 2b**). In melanoma HT-144 cells treated with MEK inhibitor mirdametinib, dose-response curves from five independent studies overlapped closely and yielded a relatively low SEE of 8.7, indicating high inter-dataset agreement. In contrast, treatment of non-small cell lung cancer NCI-H2228 cells with the antimetabolite gemcitabine revealed widely diverging responses across datasets, resulting in a substantially higher SEE of 31.1. At equivalent doses, some datasets reported near-complete inhibition, while others showed almost no activity, making it unclear whether the cell line is truly sensitive or resistant, hence complicating biomarker discovery and cross-study validation (**Suppl. Fig. 2**).

Interestingly, SEE distributions differed significantly across mechanism-of-action classes (*p* < 10^-15^ Kruskal-Wallis test; **Fig. 2c**). Targeted agents, including hormone therapies, Hedgehog, PARP and kinase inhibitors, showed relatively low SEE, whereas kinesin inhibitors, ion transport modulators, and classical cytotoxics such as antimetabolites, DNA-targeting agents, topoisomerase and mitotic inhibitors exhibited the highest SEE distributions. Consistent with this, compounds with specific mechanisms showed lower SEE, compared to broad spectrum compounds (median SEE 11.0 vs. 12.9 respectively; *p* = 8.29×10^-4^, Wilcoxon rank-sum test; **Suppl. Fig. 3a**). Notably, per-drug SEE was not associated with the number of cell lines tested (Spearman ρ = 0.04, *p* = 0.31; **Suppl. Fig. 3b**), indicating that inconsistency reflects intrinsic drug properties rather than sampling breadth.

When grouping compounds based on clinical development stage, SEE did not differ significantly across the stages (median SEE 11.2-13.4; *p* = 0.36, Kruskal-Wallis test; **Fig. 2d**), and there was no evidence of an association between SEE and stage (Spearman correlation ρ = -0.04, *p* = 0.32), indicating that inconsistency in drug response is not systematically related to clinical development phase. In contrast, SEE distributions differed significantly across datasets (*p* < 10^-15^, Kruskal-Wallis test; **Fig. 2e**). Some pharmacogenomic resources, such as FIMM and Jaaks, displayed relatively narrow SEE profiles, whereas others, including PRISM and Nair, showed broad SEE distributions with right-shifted tails extending to higher SEE values. Hierarchical clustering of pairwise differences in SEE further confirmed that the latter datasets clustered together, reflecting higher inconsistency profiles (**Fig. 2f**).

To identify the main sources of inconsistency, we correlated SEE with four broad feature groups: cell-line characteristics, drug properties, experimental parameters and dose-response curve features (**Fig. 2g**). Among these, dose-response curve features, including maximum effect, IC_50_, and slope, showed the strongest median association with SEE levels. Drug properties and experimental factors showed weak associations, whereas cell line features had the lowest contribution to inconsistency. At the individual-feature level, dose-response descriptors dominated the top correlates of SEE, led by maximum effect, activity at low concentrations, slope, Hill coefficient, and IC_50_ (**Fig. 2h**). Several experimental factors, such as assay format, readout technology, and treatment duration, as well as drug structural fingerprints, also ranked among the top correlates.

Together, these analyses indicate that cross-dataset inconsistency in drug response is driven predominantly by how responses are quantified, followed by how experiments are conducted, with drug properties exerting a secondary influence and cell line molecular features contributing only minimally. Because cell lines serve as proxies for patients in precision medicine, these findings suggest that individualized drug screens can yield consistent results, provided that analytical and experimental protocols are harmonized across laboratories and studies.

### Development and benchmarking of drug response metrics for consistency

Given the strong influence of dose-response curve features on reproducibility, we next systematically benchmarked more than 20 currently used drug response metrics, ranging from single-point readouts (e.g., IC_50_ and activity at fixed concentration) to composite metrics (e.g., area under the curve, activity area, drug sensitivity score) using the iDRR resource across all 15 integrated datasets. For each metric, we computed the Spearman correlation of the metric’s values between all pairs of datasets sharing the same drug-cell line experiment. Single-point metrics exhibited poor cross-dataset agreement, with median correlations typically below 0.4, and even the widely-used composite metrics rarely exceeded median correlation of 0.5.

To address these limitations, we developed Drug Response Score (DRS), which leverages information from the full fitted dose-response curve, while emphasizing the most informative concentration range (**Fig. 3a**). DRS integrates the drug response above a fixed activity threshold (default *T*=10%), while weighting each concentration with a hyperbolic function centered at the IC_50_, so that doses near the inflection point contribute most to the DRS values, whereas typically noisy extremes at very high or low doses contribute less. Across the integrated iDRR dataset, DRS achieved the highest cross-dataset reproducibility among all the evaluated metrics, with median inter-dataset correlations substantially higher than those of conventional metrics (*p* < 0.01, Wilcoxon signed-rank test; **Fig. 3b**). For example, comparing the Jaaks and GDSC2 datasets across their 3,848 shared experiments yielded a Spearman DRS correlation of 0.92 (*p*<0.01), whereas AUC and the other response metrics showed notably weaker consistency.

**Figure 3.**
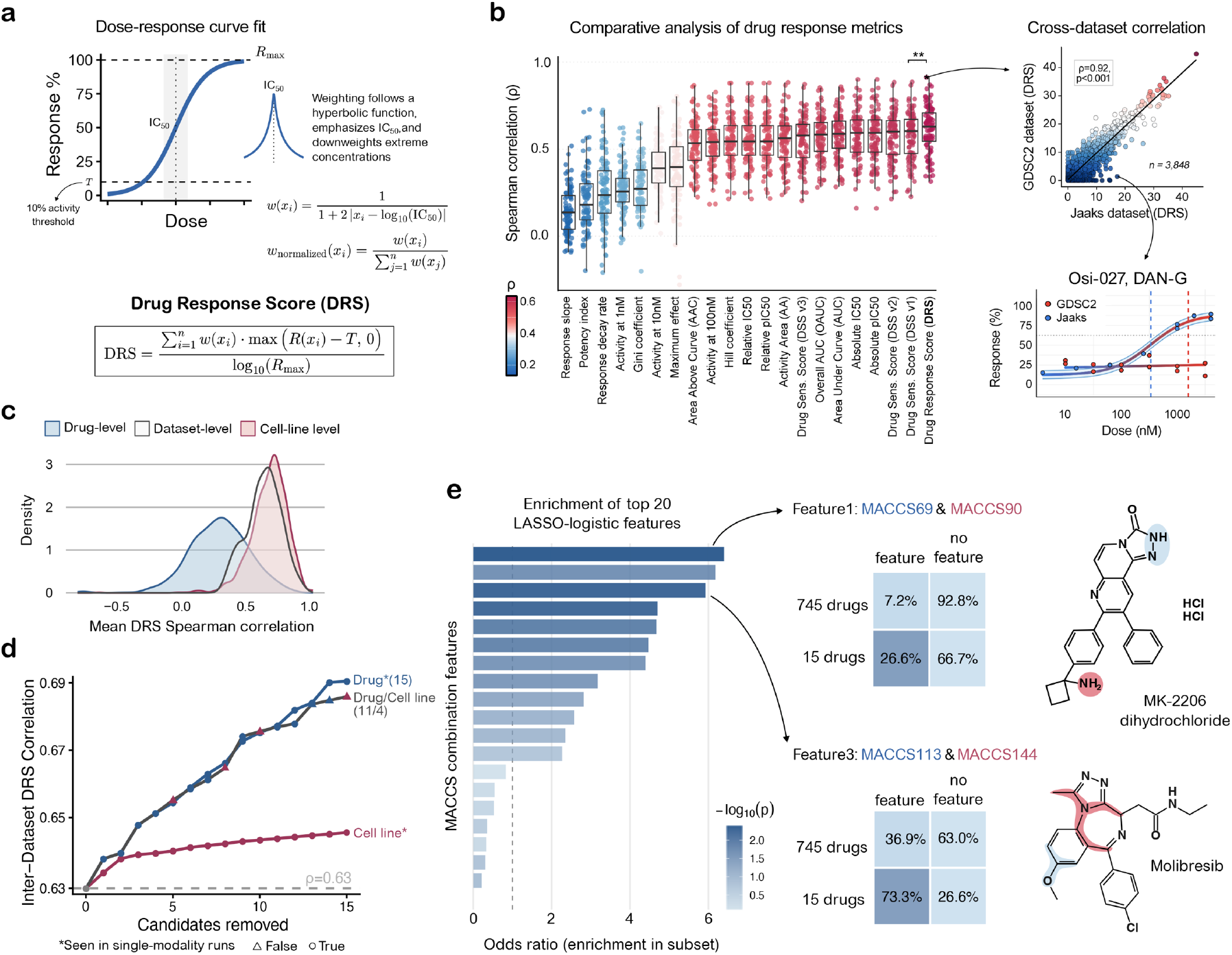
Design and evaluation of the novel Drug Response Score (DRS). (**a**) Calculation of the Drug Response Score (DRS). A 4PL curve is fit separately to each drug-cell line profile, and concentrations are weighted with a hyperbolic function centered at the IC_50_ to emphasize pharmacologically more informative doses and down weight extremes. The DRS aggregates weighted activity above a fixed threshold *T*, normalized by the maximum response. DRS ranges between 0 and 50, where 0 indicates no activity above the minimum threshold. (**b**) Benchmarking of commonly used drug-response metrics across datasets. Boxplots show the distribution of inter-dataset Spearman correlations for each metric, with the DRS (right) achieving the highest median correlation. An example scatterplot highlights strong cross-dataset concordance with DRS between the Jaaks and GDSC2 studies, and the lower panel shows a representative dose-response curve (Osi-027 in DAN-G cells). (**c**) Distributions of the average pairwise Spearman correlations of DRS between datasets, computed separately at the drug, cell line, and dataset levels. For each drug (blue), we calculated the correlation of its DRS profile across all pairs of datasets and averaged these values; the same procedure was applied to each cell line (red) and each dataset (gray). Drug-level correlations were significantly lower than cell line or dataset-level correlations (Kruskal-Wallis χ^2^(2) = 1,004, *n* = 1,937, *p* < 10^-20^, effect size H = 0.518; pairwise Wilcoxon BH-adjusted *p* < 10^−20^) and more dispersed (Levene’s F(2,1934) = 85.25, *p* < 2.2 × 10^−16^; drug-level IQR = 0.314 vs. 0.163–0.173 for dataset/cell line). Pairwise Kolmogorov-Smirnov tests confirmed differing distribution shapes (drug vs. cell line D = 0.734, adjusted *p* < 10^−20^; dataset vs. drug D = 0.664, adjusted *p* = 3.54 × 10^−32^; dataset vs. cell line D = 0.201, adjusted *p* = 1.86 × 10^−3^). (**d**) Change in inter-dataset DRS correlation as individual drugs, drug-cell line pairs, or cell lines are sequentially removed. Removing a small number of drugs or drug-cell line pairs markedly increased cross-dataset agreement, whereas removing cell lines had minimal effect. The grey dotted line indicates the baseline inter-dataset DRS correlation (ρ = 0.63). (**e**) Enrichment of top 20 pairwise MACCS substructural features among the 15 most discordant compounds. Bars show odds ratios from a LASSO-regularized logistic regression comparing the 15 discordant compounds to the remaining 745 drugs. Two enriched feature pairs are highlighted with example compounds: MK-2206 dihydrochloride (MACCS keys 69 and 90) and Molibresib (MACCS keys 113 and 144). Contingency tables show the prevalence of each feature pair in discordant versus remaining compounds.

We next asked at what level of aggregation the remaining inconsistency is most pronounced. We computed the average pairwise correlation of DRS profiles across drugs, cell lines and datasets (**Fig. 3c**), and visualized their correlation distributions. Drug-level correlations were markedly lower and more dispersed than those observed at the dataset or cell line level (*p* < 2.2×10^−16^, Kruskal-Wallis test; **Fig. 3c**), indicating that the identity of the compound is the primary source of residual cross-dataset inconsistency. To further investigate this, we used a beam search to identify the most influential compounds and examined how their removal affected global inter-dataset correlations (**Fig. 3d**). Eliminating a relatively small subset of compounds (*n*=15, <2%; **Suppl. Table 2** led to a steep increase in the overall inter-dataset correlation, whereas removing cell lines had a much smaller impact, confirming that a small number of compounds account for much of the remaining disagreement between the datasets. To characterize these 15 most discordant compounds, we performed an exploratory structural analysis using LASSO-regularized logistic regression on MACCS fingerprints, which revealed that specific pairwise combinations of substructural motifs were significantly enriched in this subset compared to the remaining drug library (**Fig. 3e**). For instance, the co-occurrence of an NH-containing heterocycle and a primary amine group, as seen in MK-2206 dihydrochloride, or a fused triazole system and a methoxy-substituted aromatic ring, as in molibresib, were among the top enriched feature pairs among the most discordant compounds.

Together, these results indicate that individual compounds are the primary source of residual variability and that certain combinations of chemical substructures may be particularly susceptible to assay- or platform-specific artefacts. Importantly, these compounds should be interpreted as candidates requiring closer experimental scrutiny rather than as definitive outliers to be excluded in all downstream analyses.

### DRS enhances reproducibility in patient-derived organoids and patient cells

To test whether DRS performance generalizes beyond immortalized cell lines, we applied the same modelling and scoring framework to two independent functional precision oncology resources that use fresh living biopsies to optimize cancer treatments for individual patients (**Fig. 4a,b**). The PMC Neuroblastoma dataset comprises patient-derived organoids (PDOs) generated from the tumors of 15 children with neuroblastoma, each screened against 200 compounds at six doses^42^. The SMARTrial is based on *ex vivo* drug sensitivity testing of primary samples from 80 patients with acute myeloid leukemia across five doses^43,44^. All dose-response measurements in these datasets were performed as technical replicates, yielding independent dose-response curves for each drug-patient pair. For each replicate, we fitted 4PL dose-response models separately and computed the full panel of drug-response metrics.

**Figure 4.**
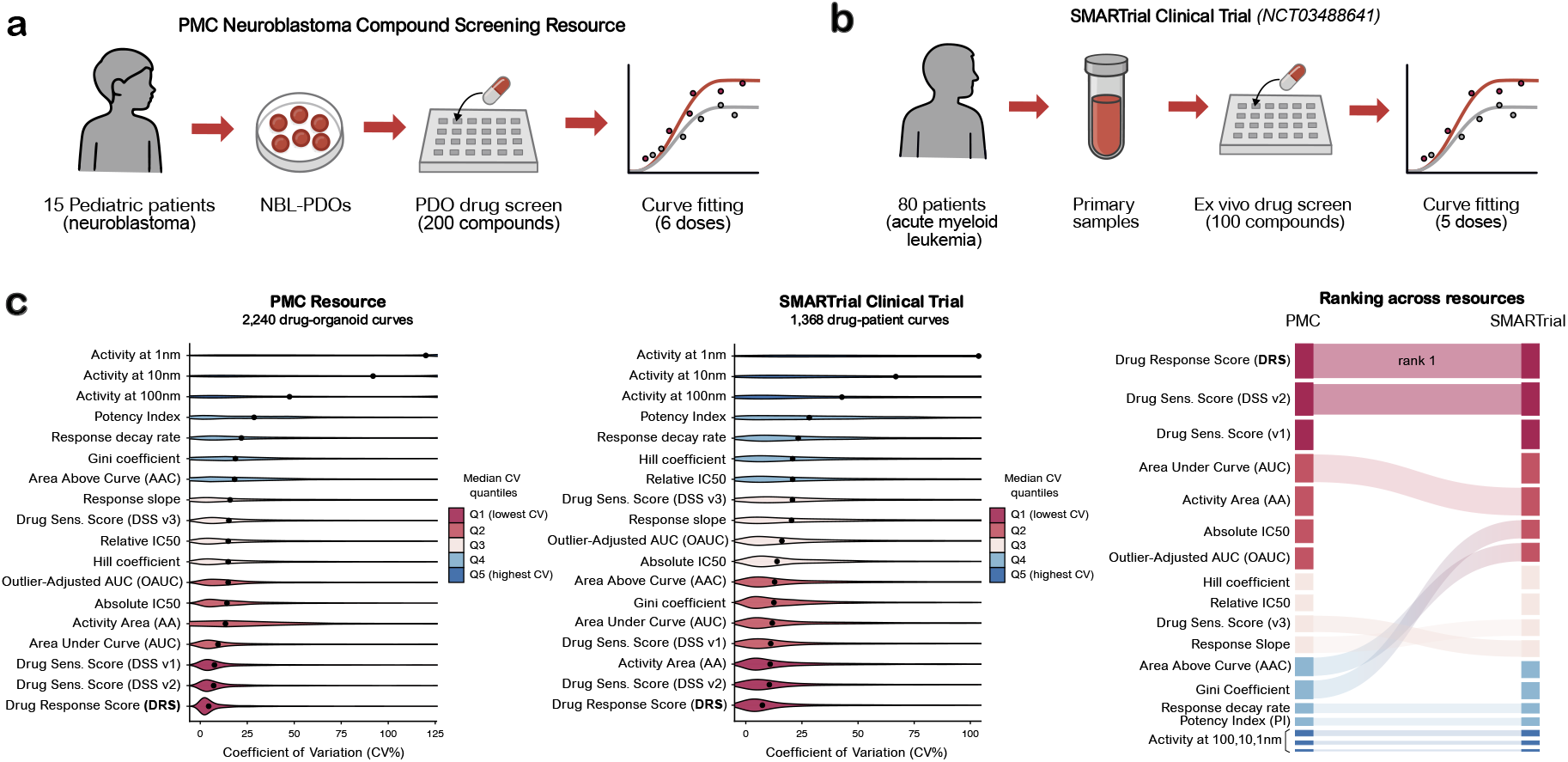
Validation of Drug Response Score in patient-derived organoids and primary patient cells. (**a**,**b**) Schematic of two functional precision oncology resources used to benchmark response metrics. (**a**) The PMC Neuroblastoma resource consists of drug screening on patient-derived neuroblastoma organoids (NBL-PDOs), generated from 15 pediatric patients, followed by multi-dose-response quantification. (**b**) The SMARTrial comprises *ex vivo* drug screening of primary samples from 80 adults with acute myeloid leukemia, with curve fitting performed on multi-dose drug-patient response profiles. Replicate dose-response curves are fitted with 4PL models and summarized using multiple drug-response metrics. (**c**) Reproducibility of drug-response metrics across the replicate measurements. For the PMC Neuroblastoma organoids (left) and SMARTrial primary samples (middle), the coefficient of variation (CV%) is shown for each metric across drug-organoid or drug-patient pairs, with the shaded regions denoting median CV quantiles. The right panel ranks metrics by their overall variability across both resources, highlighting that the Drug Response Score (DRS) consistently shows the lowest CV, and thus the highest technical reproducibility.

We quantified the assay robustness as the coefficient of variation (CV%) across replicate measurements for each response metric. In both the organoid and the primary-sample screens, DRS was consistently ranked as the top reproducible metric, exhibiting the lowest median CVs and narrower distributions across replicates, when compared to the existing single point and composite drug response metrics (**Fig. 4c**). When the metrics were ranked by their CV across both resources, DRS and its closely related DSS variants achieved the most reproducible performance, whereas the single-point metrics were less stable (**Fig. 4c, right panel**). Notably, the reproducibility of the commonly-used IC_50_ metric was in the middle of the consistency continuum.

## Discussion

Large-scale pharmacogenomic profiling has become a cornerstone of precision oncology and drug discovery, yet nominally identical experiments often yield discordant drug-response measurements across studies and datasets^15,24^. This reproducibility gap limits biomarker discovery, model development, and translation of the *in vitro* or *ex vivo* drug sensitivity into clinical practice^16,45^. Wide variability in response patterns has further led most predictive machine learning models to rely on one or two pharmacogenomics datasets, rather than systematically integrating information across multiple resources^18,19^. Here, we integrated a total of 15 monotherapy and combination screening datasets into a harmonized resource, comprising 760 compounds and 1,111 cancer cell models, resulting in nearly ten million dose-response observations. Using a unified four parameter logistic curve fitting, we defined a consensus dose-response curve for each drug-cell line pair and introduced the Standard Error of Estimate (SEE) as a quantitative measure of inconsistency between the datasets. Together, iDRR and SEE metric provide a basis for benchmarking reproducibility at a larger-scale and for systematically identifying technical and biological factors that shape cross-dataset discordance.

Existing pharmacogenomic databases, including PharmacoDB^27^ and CellMinerCDB^28^, have made important contributions by aggregating datasets, harmonizing annotations, and providing tools for cross-database exploration and biomarker analysis. However, these platforms typically integrate a subset of available monotherapy screens and focus on data access and visualization rather than on systematic assessment of reproducibility. By contrast, the iDRR resource unifies 15 large-scale pharmacogenomic datasets into a single harmonized framework comprising nearly ten million dose-response measurements, including, for the first time, single-agent readouts extracted from five drug-combination studies. Beyond aggregation, iDRR provides consensus dose-response curves for each drug–cell line pair, per-pair inconsistency estimates (SEE), and an optimized response metric (DRS) specifically developed to maximize cross-dataset concordance. These analytical capabilities are not available in existing resources. Our analyses demonstrate that inconsistency is driven predominantly by how drug responses are quantified and how experiments are conducted, rather than by intrinsic cell line heterogeneity. Dose-response characteristics showed the strongest association with higher inconsistency, followed by experimental parameters, such as treatment duration, plate format and viability readout. Among the experimental factors, the highest positive correlates were the PRISM dataset indicator and its specific assay components, namely Luminex readout and 384-well plate format, whereas the other resources, for example GDSC1, which uses resazurin-based readout and 96 well plate assay, which associated with lower SEE. These associations do not prove that specific assay formats are more inconsistent than others, rather they highlight that the choice of readout and plate format can have severe effects on reproducibility when looking at inter-dataset agreement. Structural fingerprints emerged as the most informative drug features, yet remained much smaller in contribution than that of metric-related or experimental factors.

Notably, we observed marked differences in inconsistency across drug classes. Hormone therapies, PARP inhibitors and many kinase inhibitors exhibited relatively low SEE, whereas antimetabolites, DNA-targeting agents, and topoisomerase and mitotic inhibitors were substantially more variable between the datasets. The lack of a systematic relationship between SEE and clinical phase indicates that inconsistency is not simply a property of drug candidates and probes, but it persists for agents at all stages of clinical development. Together, these results argue that reproducibility should be evaluated at the level of individual compounds and mechanisms, not assumed from clinical maturity or overall assay quality.

Previous works have shown that with rigorous quality control (QC) measures and accurate response metrics, discordance in large pharmacogenomic screens can be reduced^35,46,47^. Because QC acts upstream of response modelling and metric selection^48,49^, an effective filtering of low-quality experiments is often the most immediate and impactful step for not only improving cross-dataset agreement, but also when building more consistent large-scale pharmacogenomics resources. Plate-level artifacts and other systematic technical biases, including edge and evaporation effects, plate row and column gradients from dispensing, and temperature-related drift, can inflate variance and distort downstream dose-response curves. These technical artifacts are a primary source of inconsistency observed both within and between studies. We recently showed that control-independent, curve-fitting residual metrics, such as the Normalized Residual Fit Error (NRFE)^50^, are highly effective at flagging compromised experiments, filtering of which significantly improves cross-study correlation. Therefore, NRFE-style QC metrics can serve as the first-line filter followed by SEE to quantify remaining inconsistency between technical replicates and benchmark new datasets against existing resources to pinpoint potentially problematic dose-response curves.

Given the strong influence of response metrics on the cross-dataset concordance, we systematically benchmarked more than 20 commonly used drug-response measures. Single-point readouts, such as activity at fixed concentrations and absolute or relative IC_50_, showed a relatively poor agreement between datasets, and even composite metrics, such as AUC, AAC, and DSS variants, achieved only modest median correlations. To address this limitation, we developed the Drug Response Score (DRS), which leverages the information from a full fitted dose-response curve, while emphasizing pharmacologically informative concentrations near the IC_50_^21^. By down-weighting noisy extremes at very low or high doses, DRS achieved substantially higher inter-dataset correlations than the existing metrics and it was robust across a wide range of compounds, cell lines, and datasets. These improvements were not confined to immortalized cell lines only; in independent patient-derived neuroblastoma organoids and primary AML samples, DRS consistently exhibited the lowest variation across replicates, outperforming traditional metrics in both personalized medicine settings.

These findings help to explain a recurring pattern seen in literature, where many predictive machine learning models developed for drug-response prediction are based on a single major resource, such as GDSC or CCLE, rather than being trained and validated across integrated datasets^51,52^. When the drug response labels are inconsistent across studies, cross-resource generalization becomes limited, with ML models and predictions reflecting dataset-specific idiosyncrasies, rather than true biological signals. By quantifying inconsistency at the level of drug-cell line experiments, SEE provides a practical way to contextualize measurements across resources, particularly useful when creating new large-scale high-quality resources. Our results suggest that a large part of the inconsistency is metric-dependent^23,53^, motivating the development of DRS. In ML modelling, DRS offers a more consistent response outcome, providing a reliable target for multi-resource biomarker discovery and predictive modelling.

The harmonized iDRR resource of 15 datasets and the DRS metric have several practical implications for the design and analysis of future pharmacogenomic studies. First, SEE provides an intuitive benchmark to identify compounds, cell lines, and datasets that contribute disproportionately to inconsistency. Our beam-search analysis showed that a relatively small subset of compounds accounts for a large fraction of the remaining discordance, suggesting that targeted follow-up on these problematic agents, for example re-testing under more standardized conditions or investigating assay interference, could significantly improve global reproducibility. Second, DRS consistently outperformed all conventional metrics used for drug response quantification in cross-dataset reproducibility, making it a strong candidate as a standard metric for pharmacogenomic analyses, including biomarker discovery and machine-learning modelling, where robust and reproducible drug-response quantification is essential. Third, the integration of monotherapy screens with single-agent readouts from combination experiments illustrates how diverse resources can be pooled into a coherent framework to maintain transparent and comparable readouts across heterogeneous studies.

This study has some important limitations. Although our analysis spans a large number of compounds, assays and cell models, it remains confined to *in vitro* and *ex vivo* systems and cell viability readouts. Many sources of variability that affect clinical treatment response, such as pharmacokinetics, tumor microenvironment, and host factors, are not captured in cell lines, organoids, or short-term *ex vivo* cultures. Moreover, despite our efforts to harmonize identifiers and modelling approaches, residual heterogeneity in experimental protocols, compound handling, and data processing is unavoidable and may influence SEE estimates. The beam-search approach to identifying discordant candidates is heuristic and may depend on the specific set of datasets and correlation thresholds considered; the list of problematic compounds should therefore be interpreted as a practical guide, rather than a definitive classification. Finally, while we demonstrate that DRS improves technical reproducibility, systematic evaluation of its ability to predict patient benefit will require larger patient cohorts with matched *ex vivo* screening and clinical outcome data.

Despite these caveats, our work provides a quantitative and extendable framework for assessing and improving reproducibility in pharmacogenomic screens. By jointly modelling dose-response measurements across 15 major resources, we show that cross-dataset agreement is shaped primarily by adjustable factors (i.e., how responses are quantified and experiments are conducted), and that pharmacologically informed weighting of the dose-response curve can substantially enhance concordance. The publicly available iDRR resource, together with the open-source implementation of DRS metric, offers the community a unified platform for methods development, benchmarking, dataset curation, and reproducible biomarker discovery. As pharmacogenomic profiling continues to expand into new model systems, phenotypic readouts, and clinical workflows, systematic evaluation of reproducibility will remain essential to realizing the promise of precision oncology.

## Methods

### Data sources and integration

We assembled the largest pharmacogenomic resource to date by curating single-agent drug-response data from ten major initiatives (**Table 1; Suppl. Table 1**): CCLE^1^, CTRPv2^4^, gCSI^24,31^, GDSC1^3,30^, GDSC2^2,3,30^, GRAY^32,33^, FIMM^34,35^, PRISM^5^, NCI60^29,36^, and UHNBreast^54,55^. We also incorporated five drug-combination studies: AstraZeneca^37^, Bashi et al.^38^, Jaaks et al.^39^, Nair et al.^40^, and O’Neil et al.^41^; for these, single-agent responses were extracted from the dose-combination matrix edges, where one compound was titrated while the partner compound was fixed at zero concentration. To standardize drug identifiers, all compounds were harmonized using PubChem compound identifiers and all cell lines using Cellosaurus accession identifiers. To ensure statistical robustness, only compound-cell line pairs measured in at least two datasets were retained, yielding 195,317 unique pairs spanning 760 compounds and 1,111 cell lines.

### Cell line metadata and compound features

To systematically evaluate whether chemical, biological, or experimental factors contributed to variability in drug-response measurements, we curated and harmonized features at three levels: compounds, cell lines, and dataset-specific experimental conditions.

#### Molecular feature extraction

We retrieved SMILES strings for each compound from PubChem using its API and encoded structural information with seven fingerprinting methods: Extended Connectivity Fingerprints (ECFP4 and ECFP6), Molecular ACCess System keys (MACCS), Electrotopological State (EState), PubChem fingerprints, Klekota-Roth descriptors, and Extended 3D Fingerprints (E3FP). Physicochemical descriptors included molecular weight, calculated octanol-water partition coefficient (ALogP), hydrogen bond donor and acceptor counts, polar surface area, rotatable bond count, aromatic ring count, heavy atom count, and Lipinski’s Rule-of-Five violations. Mechanism of action and clinical development phase annotations were obtained from PubChem^56^ and DrugBank^57^, and manually curated.

#### Cell line feature extraction

Cell line metadata were curated from Cellosaurus^58^ including patient demographics (sex, age at sampling), disease classification, and anatomical site of origin. Cellosaurus accessions were cross-referenced with DepMap^59^ to obtain microsatellite instability status, treatment history, and ploidy. Metastatic status was validated through triangulation of DepMap, Cell Model Passports^60^, and site-disease concordance. Multi-omic features of cell lines were retrieved from DepMap, including transcriptomics (RNA-seq), chromatin accessibility (ATAC-seq), copy-number alterations, mass-spectrometry (MS)-based proteomics, and reverse-phase protein arrays (RPPA).

#### Dataset-specific feature extraction

For each pharmacogenomic dataset, we collected experimental parameters from the original publications and supplementary materials, including publication year, cell line culture conditions (medium and serum content), assay plate format (96-, 384-, or 1536-well), treatment duration (48-120h), and viability readout (CellTiter-Glo, Luminex, Resazurin/SYTO60, Sulforhodamine B, or SYTOX Green).

### Consensus dose-response modelling

To obtain standardized response estimates across studies, we implemented a consensus curve-fitting approach. For each drug-cell line pair measured in ≥2 datasets, all available dose-response points were pooled and fitted with a four-parameter logistic regression (4PL):

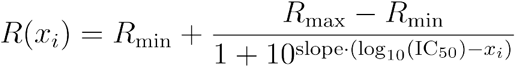

where *R*(*x*_*i*_) is the response (percent inhibition) at log concentration *x*_*i*_, *R*_*min*_ is minimum response (typically 0), *R*_*max*_ is the maximum response parameter from the fitted curve, slope is the Hill slope parameter and *IC*_50_ is the half maximal inhibitory concentration. The pooled 4PL fit provides a unified, dataset-independent estimate of drug sensitivity that reduces bias compared with study-specific curve fitting.

### Benchmarking of drug-response metrics

To assess the reproducibility of commonly used drug-response metrics, we benchmarked a comprehensive panel of conventional metrics reported in the literature. These included single-point metrics, such as IC_50_ (absolute, relative, pIC_50_), response slope, Hill coefficient, and drug activity at fixed concentrations (10 nM, 100 nM, 1 nM), as well as composite scores that integrate the information from a full dose-response curve, including area under the curve (AUC), outlier-adjusted AUC (OAUC), activity area (AA), area above the curve (AAC), Gini coefficient (a measure of inequality in the distribution of response values across tested concentrations)^61^, response decay rate, and three versions of the Drug Sensitivity Score (DSS v1-v3)^22,49^. An overview of these metrics is provided in **Suppl. Fig. 4**. For each metric, reproducibility was assessed by calculating Spearman correlation across pairwise dataset comparisons of overlapping drug-cell line measurements.

### Development of Drug Response Score (DRS)

To overcome limitations of existing drug-response metrics, we developed the Drug Response Score (DRS), a proximity-weighted metric that emphasizes pharmacologically relevant dose ranges around the IC_50_ region. The DRS is formally defined as:

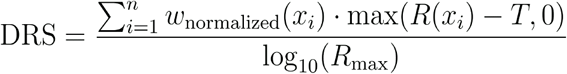

where are log-transformed concentration points sampled uniformly in log space between *log*_10_(*C*_*min*_)and *log*_10_(*C*_*max*_), (*C*_*min*_) is the minimum concentration, (*C*_*max*_) is the maximum concentration, *n* is the number of sampling points (100 by default), *R*(*x*_*i*_) is the fitted response (percent inhibition) at log-concentration *x*_*i*_, *T* is the minimum activity threshold (10% by default), *R*_*max*_ is the maximum response parameter from the fitted 4PL curve, and *w*_normalized_(*x*_*i*_) is the normalized weight function at point *x*_*i*_.

The weights were defined to decay hyperbolically with distance from the IC_50_:

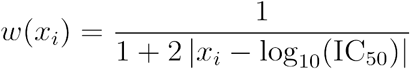

and normalized such that ∑w(x_i_) = 1. This hyperbolic form was chosen because it sharply emphasizes doses near the IC_50_, where most pharmacological importance lies, while still allowing information from the broader dose ranges to contribute to the DRS calculation. The threshold *T* = 10% reflects the minimal level of response commonly used to distinguish background noise from biologically meaningful activity and the choice of *N* = 100 sampling points provides a stable approximation to continuous integral of the weighted response without imposing substantial computational cost. DRS ranges between 0 and 50, where 0 indicates no activity above the minimum threshold.

### Evaluating inconsistency

To assess consistency, we defined a Standard Error of Estimate (SEE) to quantify deviations of observed responses from the consensus 4PL fit across datasets:

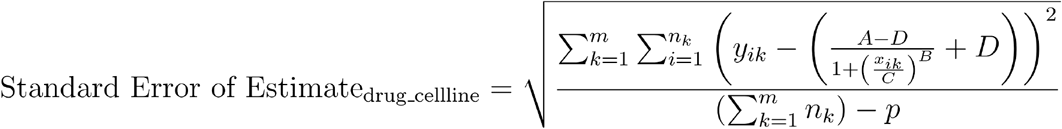

Here, *m* is the number of datasets, *y*_*ik*_ is the observed response for the ith observation in dataset *k, x*_*ik*_ is the dose for the ith observation in dataset *k, n*_*k*_ is the number of observations in dataset *k*, [*A,B,C* and *D*] are the parameters of the 4 parameter logistic model, *p* is the number of parameters in the model (4 for the 4PL method). Lower SEE values reflect reproducible responses, while higher values indicate greater cross-study inconsistency. The minimum SEE is 0 and the theoretical maximum is 100, corresponding to the full range of the viability scale. This single interpretable score provides a robust benchmark for evaluating drug-response reproducibility across datasets.

### Systematic analysis of factors contributing to inconsistency

To determine the factors that influence reproducibility of drug-response measurements, we assembled a comprehensive feature set spanning multiple domains. For each drug-cell line pair, we included (i) dose-response curve characteristics derived from 4PL fits (Hill slope, maximum effect, activity at fixed concentrations); (ii) cell line characteristics and multi-omic features (sex, age at sampling, anatomical site, disease classification, metastatic status, microsatellite instability, ploidy, prior treatment history, transcriptomic expression, chromatin accessibility, copy-number alterations, proteomics, and RPPA profiles); (iii) dataset-specific experimental parameters (culture medium, serum concentration, assay format, treatment duration, and viability readout) and binary dataset indicators denoting whether a given drug-cell line pair was tested in a particular dataset; (iv) compound descriptors (structural fingerprints, physicochemical properties, mechanism of action, and clinical development stage).

To harmonise heterogeneous feature types, we transformed all variables into common numeric representation prior to association testing: binary attributes were encoded as 0/1, categorical metadata were one-hot encoded and continuous variables were transformed as needed. For each feature *x*, we computed Spearman’s rank correlation (ρ) between *x* and SEE across all drug-cell line pairs. Two-sided p-values for the null hypothesis of zero correlation were adjusted for multiple testing with Benjamin Hochberg false discovery rate (FDR) procedure for each feature as well as for each feature set. Given the large number of drug-cell line pairs, even modest correlations were statistically significant after FDR correction; we therefore focused on the absolute correlation magnitude (|ρ|). For interpretability, correlations were classified as weak (|ρ| < 0.10), moderate (0.10 ≤ |ρ| < 0.30) and strong (|ρ| ≥ 0.30). Features that were weakly correlated with SEE were interpreted as relatively robust to cross-study variability, whereas those strongly correlated with SEE were considered potential contributors to inconsistency.

### Beam-search optimization search to identify discordant compounds and cell lines

To identify the minimal set of compounds whose removal maximizes cross-dataset correlation, we implemented a heuristic beam search^62,63^ on DRS measurements across 195,317 drug-cell line curves. Starting from a full dataset (no candidates removed), we iteratively constructed new states by removing one candidate from the total set (PubChem CIDs for drugs and Cellosaurus accession identifiers for cell lines) and recomputing the Spearman correlation after excluding all rows corresponding to the experimental drug-cell line DRS values associated with those identifiers (**Suppl. Fig. 5**). At each depth (i.e., number of candidates removed), we employed a beam of 3, meaning all states generated from the beams were evaluated and ranked by their updated correlation, but only the top three states were retained for further expansion. We performed separate searches that allowed removal of (i) drugs only, (ii) cell lines only, and (iii) both drug and cell lines, and stopped when the global DRS correlation reached a target of 0.7 or when no further improvement was observed. This procedure resulted in one parsimonious set of candidates for each scenario; because it is heuristic and depends on the chosen correlation target and beam width, the resulting subsets should be interpreted as practical flags rather than definitive outliers.

### Structural pattern analysis of discordant compounds

To investigate whether structural features distinguish the 15 compounds whose removal most improved inter-dataset correlation, we used a LASSO-regularized logistic regression on MACCS fingerprints. Each compound in the 760-compound dataset was encoded as a 166-bit MACCS key vector (RDkit v2022.09.5), and compounds in the predefined subset were assigned the label 1, while all others were assigned the label 0. To capture co-occurrence patterns, we generated second-order interaction features corresponding to pairwise logical AND combinations of MACCS bits. A logistic regression model with an L1 penalty was fit to this structural feature matrix. Features with non-zero coefficients in the fitted model were interpreted as MACCS combination keys that were most informative for distinguishing compounds in the subset from the rest of the library, and thus as candidate structural motifs associated with inconsistent behavior.

### Application of drug response scores to patient-derived cells

We applied the identical analysis pipeline to two independent patient-derived dose-response resources: (i) the PMC Neuroblastoma Compound Screening Resource (accessed October 20, 2025)^42^ and (ii) the SMARTrial (NCT03488641) clinical trial^44,64^. For each drug-sample pair, we prioritized the technical replicate pair R1-R2; if R1 was unavailable but R3 existed, we used R2-R3; if R2 was unavailable but R3 existed, we used R1-R3. Pairs with < 3 unique dose levels were excluded from model fitting.

Each available replicate experiment was fitted with a four-parameter log-logistic (4PL) model as described earlier. From each fitted curve, we computed standard pharmacologic metrics, including IC_50_ (absolute and relative), response slope, Hill coefficient, activity at fixed concentrations (10 nM, 100 nM, 1 nM), area under the curve (AUC), outlier-adjusted AUC (OAUC), activity area (AA), area above the curve (AAC), Gini coefficient, response decay rate, three versions of the Drug Sensitivity Score (DSS v1-v3) and Drug Response Score (DRS).

Reproducibility was summarized per metric by the coefficient of variation (CV) across the replicates. Let *k* index the *x* metrics and *r* index replicates; *m*_*r,k*_ denotes the value of metric *k* in replicate *r*. We report *CV*% where lower *CV*% indicates better reproducibility.

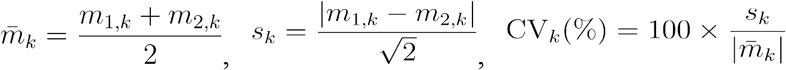

## Code and Data Availability

The integrated and harmonized drug response resource (iDRR), including consensus drug-response curves, standard error of estimate (SEE) values, Drug Response Score (DRS) is publicly available. The complete resource has been deposited at Zenodo (https://doi.org/10.5281/zenodo.18775863) and is distributed as a fully documented R package (https://github.com/kris-nader/idrrR). An interactive web portal enables users to explore drug-response metrics and dose-response relationships across all harmonized measurements (https://aittokallio.group/iDRR/).

Raw and processed pharmacogenomic data for the large-scale screens (monotherapy and combination studies) are available from the original sources as described in the corresponding publications and summarized in **Supplementary Table 1**. Single-agent drug-response data from the AstraZenica-Sanger Drug Combination Prediction DREAM Challenge are available via Synapse (syn4231880), subject to the associated access terms. The cell line multi-omics data were obtained from the DepMap Public 24Q2 release (https://plus.figshare.com/articles/dataset/DepMap_24Q2_Public/25880521/1).

Drug-response data for the PMC Neuroblastoma Compound Screening Resource can be interactively explored and downloaded through the R2: Genomics Analysis and Visualization Platform at the PMC Neuroblastoma Compound Screening Resource portal (https://hgserver1.amc.nl/cgi-bin/r2/main.cgi). Data from the SMARTrial cohort, including *ex vivo* drug-response profiles and accompanying analysis scripts, are publicly available at the SMARTrial GitHub repository (https://github.com/PeterBruch/SMARTrial).

## Acknowledgements

We thank the DDCB core facility (FIMM HTB unit) supported by the University of Helsinki and Biocenter Finland. Funding support: AI: Ida Montin Foundation grant. TA: Research Council of Finland (grants 340141, 344698, 367855); the Cancer Society of Finland, the Norwegian Cancer Society (grants 216104 and 273810), Norwegian Health Authority South-East (grants 2020026 and 2023105), the Sigrid Jusélius Foundation, and iCAN – Digital Precision Cancer Medicine Flagship (iCAN-MULTIDRUG). This work was supported by the REMEDi4ALL project, which has received funding from the European Union’s Horizon Europe research and innovation programme under grant agreement No 101057442. Views and opinions expressed are those of the author(s) only and do not necessarily reflect those of the European Union, who cannot be held responsible for them.

