## Supplemental Table for "Improving reproducibility in pharmacogenomic screens through cross-study benchmarking"

**Supplement Table 1. Overview of pharmacogenomic datasets integrated and harmonized in iDRR.**

| Type | Dataset | Key References | Source/Description |
|---|---|---|---|
| Monotherapy | CCLE | Barretina et al Nature 2012 | Cancer Cell Line Encyclopedia. Broad Institute |
| | CTRPv2 | Seashore-Ludlow et al. Cancer Discov 2015 | Cancer Therapeutics Response Portal. Broad Institute. |
| | FIMM | Pemovska et al. Nature 2013, Mpindi et al., Nature 2016 | Institute for Molecular Medicine Finland |
| | gCSI | Klijn et al., Nat Biotechnol 2015; Haverty et al., Nature 2016 | Genentech Cell Line Screening Initiative |
| | GDSC1 | Garnett et al., Nature 2012 | Genomics of Drug Sensitivity in Cancer (v1). Sanger Institute |
| | GDSC2 | Iorio et al., Cell 2016 | Genomics of Drug Sensitivity in Cancer (v2). Sanger Institute |
| | GRAY | Daemen et al., Genome Biol 2013; Heiser et al., PNAS 2011 | Gray Lab (OHSU) Breast Cancer Screen |
| | NCI60 | Shoemaker, Nat Rev Cancer 2006; Shankavaram et al., BMC Genomics 2009 | NCI-60 Human Tumor Cell lines screen. NCI DTP |
| | PRISM | Corsello et al., Nat Cancer 2020 | Profiling Relative Inhibition Simultaneously in Mixture. Broad Institute |
| | UHNBreast | Mammoliti et al., Sci Data 2019; Marcotte et al., Cell 2016 | University Health Network Breast Cancer Screen |
| Combination | AstraZeneca | Menden et al., Nat Commun 2019 | AstraZeneca-Sanger Drug Combination Prediction DREAM Challenge |
| | GDSC2comb | Bashi et al., Cancer Discov 2024 | GDSC Drug Combination Screen |
| | Jaaks | Jaaks et al., Nature 2022 | GDSC Drug Combination Screen |
| | Nair | Nair et al., Nat Commun 2023 | Large-scale lung cancer combination screen |
| | O’Neil | O’Neil et al., Mol Cancer Ther 2016 | Merck & Co Combination Screen |

Datasets are grouped by screen type (monotherapy vs combination). For each dataset, we report the primary references describing the screen and a brief source/description indicating the research consortium or institute.

**Supplement Table 2. Compounds pruned by beam search to enhance cross-dataset concordance.**

| PubChem CID | Drug Name | Mechanism of Action | Max Phase | Median SEE | Number of datasets |
|---|---|---|---|---|---|
| 5330286 | Palbociclib | Kinase Inhibitors | 4 | 9.55 | 10 |
| 10127622 | Selumetinib | Kinase Inhibitors | 4 | 9.97 | 12 |
| 60838 | Irinotecan | Topoisomerase Inhibitors | 4 | 17.43 | 8 |
| 23725625 | Olaparib | PARP inhibitor | 4 | 7.55 | 11 |
| 32874 | Doxorubicin | Topoisomerase Inhibitors | 4 | 16.275 | 10 |
| 24180719 | PLX-4720 | Kinase Inhibitors | 0 | 10.15 | 7 |
| 9444 | Azacitidine | Differentiating/Epigenetic Modifiers | 4 | 14.67 | 5 |
| 148124 | Docetaxel | Mitotic Inhibitors | 4 | 14.03 | 10 |
| 3385 | 5-Fluorouracil | Antimetabolites | 4 | 11.555 | 12 |
| 126941 | Methotrexate | Antimetabolites | 4 | 18.63 | 8 |
| 176871 | Erlotinib Hydrochloride | Kinase Inhibitors | 4 | 7.8 | 13 |
| 46943432 | Molibresib | Differentiating/Epigenetic Modifiers | 2 | 9.86 | 5 |
| 46930998 | MK-2206 dihydrochloride | Kinase Inhibitors | 2 | 11.35 | 11 |
| 44224160 | Osi-027 | Kinase Inhibitors | 1 | 16.715 | 7 |
| 3503 | Gossypol | Apoptotic Modulators | 1 | 19.625 | 3 |
