## Supplemental Figure for "Improving reproducibility in pharmacogenomic screens through cross-study benchmarking"

**Supplementary Information**

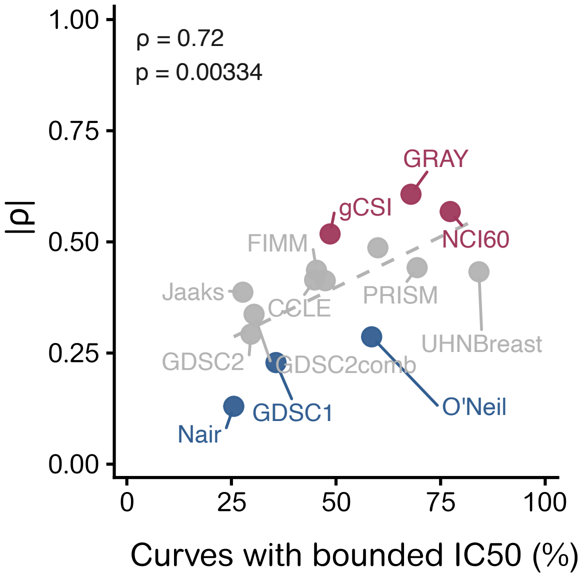


**Supplement Figure 1| Dataset-level correlation between curve quality and reproducibility.** The proportion of dose-response curves with bounded $IC_{50}$ values (indicating properly captured dose ranges) was strongly correlated with the global dose-viability correlation across pharmacogenomic datasets (Spearman rho =0.72 p=0.00334). Datasets from more focused screens (e.g., GRAY) exhibited higher quality curve fits and stronger correlations compared to discovery screens (e.g., Nair et al). Points are colored to highlight datasets with the highest (red) and lowest (blue) global Spearman correlations.

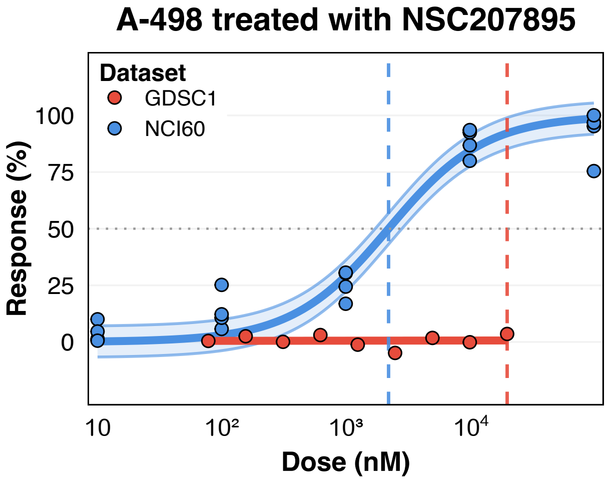


**Supplement Figure 2 | Example of cross-dataset disagreement in the dose-response profiles.** Observed responses (points) for the same drug-cell line pair (A-498 cell line treated with NSC207895) measured in two pharmacologic screens: GDSC1 (red) and NCI60 (blue) are shown across concentrations (x-axis, dose in nM on $log_{10}$ scale). Solid lines indicate the fitted 4-parameter logistic (4PL) curves. Horizontal dotted line marks the 50% response level and vertical coloured lines indicate the corresponding $IC_{50}$ estimates in the two studies. While NCI60 assay exhibited a clear sigmoidal response with increasing inhibition at higher doses, GDSC1 assay showed inactivity in the tested dose range.

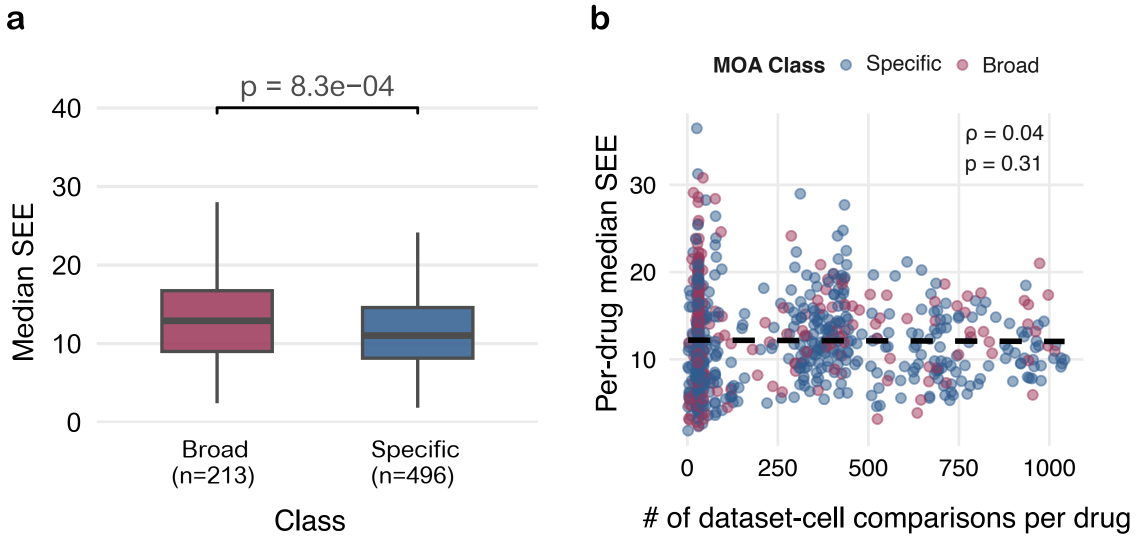


**Supplement Figure 3| Mechanism of action and Standard Error of Estimate.** (a) Boxplot of median Standard error of estimate (SEE) for compounds classified as broadly-active (n=213) and target-specific (n=496). Target-specific agents showed significantly lower SEEs compared to broad-spectrum compounds ($p=8.3 \times 10^{-4}$, Wilcoxon rank-sum test). (b) Scatter plot of per-drug median SEE against number of dataset-cell line pairs tested. No significant correlation was observed. Points are colored by MoA class (Blue, specific; red, broad).

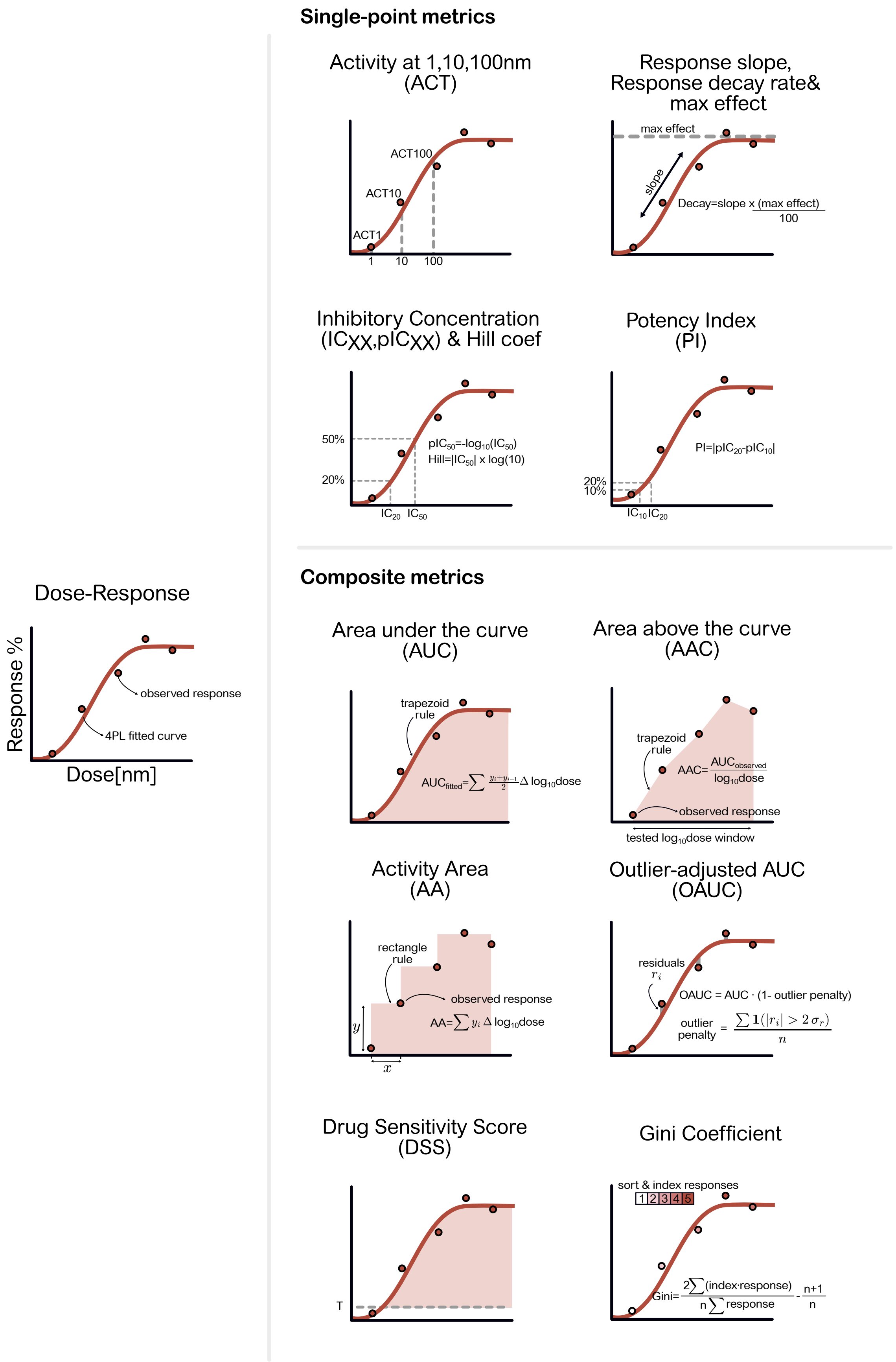


**Supplement Figure 4 | Overview of conventional dose-response metrics.** Schematic illustration how commonly used drug-response metrics are derived from a dose-response experiment. Left, observed responses (points) measured across concentrations (x-axis, dose in nM; typically analyzed on a $\log_{10}$ scale) are summarised with a four-parameter logistic (4PL) curve fitting (red). Upper panel, single-point metrics include activity at fixed concentrations (e.g., ACT1, ACT10, ACT100), inhibitory concentration, such as $IC_{50}$ and $pIC_{50}$, and curve-shape descriptors including hill coefficient, max effect, response slope, decay rate and potency index. Lower panel, composite metrics that integrate activity information over the tested dose range.

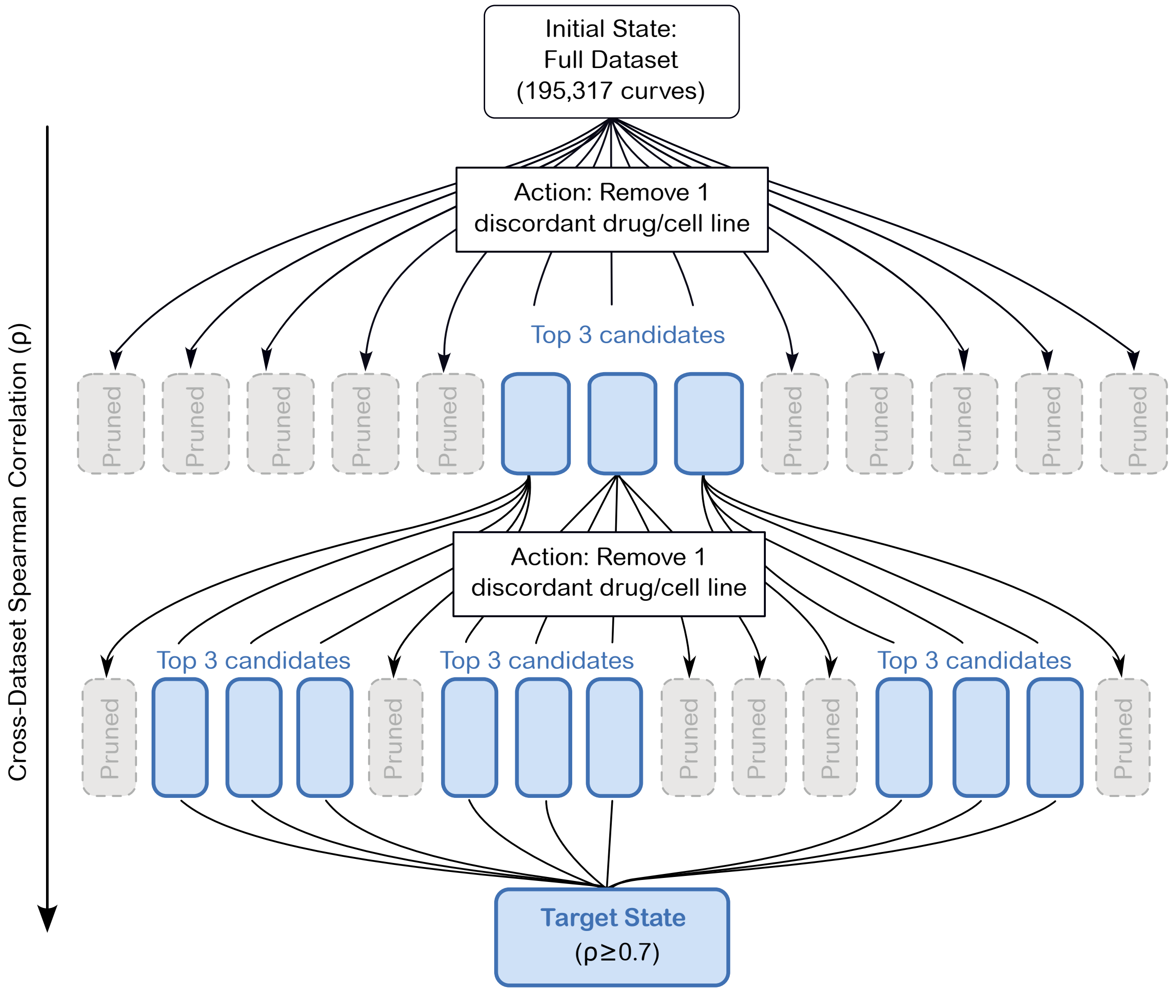


**Supplement Figure 5 | Beam-search strategy for identifying discordant compounds/cell lines to improve cross-dataset correlation.** Schematic illustrating the iterative beam-search procedure to identify a subset of drugs, cell lines or drug/cell line pairs whose removal maximizes cross-dataset correlation. The heuristic search begins with a full dataset (top). In each iteration, the algorithm generates potential states by removing either a drug or a cell line. All candidates are evaluated but only the top $k$ states ($k$=3 shown in blue) are selected for expansion as their removal leads to the most improvement in cross-dataset correlation. Suboptimal paths (dashed gray boxes) are automatically pruned. This process is repeated until the target correlation (here, 0.7) is achieved or no further improvement is found.
